# Benchmarking long-read simulators against Oxford Nanopore whole-genome sequencing data

**DOI:** 10.64898/2026.05.06.723380

**Authors:** Mona L Taouk, Danielle J Ingle, Ryan R Wick

## Abstract

**Background:** Oxford Nanopore Technologies (ONT) sequencing is increasingly used for whole-genome sequencing (WGS) across a wide range of applications. However, the platform has evolved rapidly through updates to flow cell chemistry and basecalling algorithms, altering the characteristics of the resulting sequencing data. Read simulators provide synthetic datasets with known ground truth, enabling controlled development and evaluation of methods. However, many existing simulators were developed for earlier versions of ONT sequencing or use generic long-read assumptions, and their realism for contemporary ONT data is unclear.

**Results:** We benchmarked six ONT-compatible read simulators (Badread, LongISLND, lrsim, NanoSim, PBSIM3 and SimLoRD) using a microbial genome reference and ONT R10.4.1 reads as the empirical standard. Each tool was configured to maximise realism, including training on empirical reads when supported. We compared simulated and real datasets with respect to read length, read accuracy, FASTQ quality scores and sequence error profiles. No simulator reproduced all metrics of the real data well. PBSIM3 most closely reproduced read length, read accuracy and FASTQ quality scores, making it a strong simulator for broad read-level realism. However, it did not capture important features of the real error profile, including context-dependent substitution rates and homopolymer-length errors. Badread and LongISLND better reproduced some aspects of the error profile, but showed other departures from the real data.

**Conclusion:** PBSIM3 is a good general-purpose choice for many ONT WGS simulation tasks because it reproduced several key read-level properties well. However, Badread or LongISLND may be preferable for applications where error structure is more important. No evaluated tool was realistic across all tested metrics, highlighting a gap for improved long-read simulators.

## 1. Background

Oxford Nanopore Technologies (ONT) sequencing has transformed long-read genomics by enabling real-time, portable and increasingly high-throughput sequencing ^1^. Over the past decade, ONT platforms have changed rapidly, particularly through updates to nanopore chemistry and basecalling models ^2,3^. Consequently, the characteristics of ONT data, such as error profiles and base quality scores (Q-scores), have shifted substantially over relatively short time frames of a few years ^4,5^. ONT benchmarking studies can age quickly, because their conclusions depend on the chemistries, protocols and basecalling models available at the time ^6^. Although ONT continues to evolve, recent changes in read accuracy are now more gradual than the major advances seen earlier in the platform’s development. This increasing stability is making ONT more attractive for routine, high-throughput applications where analytical pipelines must withstand formal validation and accreditation ^7,8^.

As ONT sequencing matures, its use is expanding both in high-income countries, where laboratories are now more willing to invest in validation of ONT-based workflows, and in low- and middle-income settings, where ONT’s low capital cost and portability make it a compelling alternative to some short-read sequencers ^9–11^. In both contexts, there is a need for robust, realistic benchmarking of long-read analysis tools for tasks such as read mapping, variant calling and genome assembly ^12–14^. Benchmarking using real ONT datasets is essential, but it is also constrained by sample availability, cost and the difficulty of establishing a ground truth, particularly for complex genomes ^12^. Read simulators therefore play an important role: they provide synthetic datasets where the underlying sequence is exactly known, enabling controlled, reproducible evaluation of methods across a range of experimental designs. They can also supply the large labelled training sets needed for machine learning methods such as deep neural networks ^15^.

However, many widely used ONT-compatible simulators were originally developed for earlier generations of ONT (pre-R10.4.1 flowcells) or PacBio long-read sequencing, and they may incorporate assumptions that no longer hold for modern ONT ^16–20^. For example, simulators often assumed high error rates and a simple error structure ^16,18^. Since then, improvements in ONT chemistry and basecalling have improved read accuracy, altered the balance between substitutions and indels, and changed the relationship between FASTQ Q-scores and error rates ^8^. It is therefore unclear whether existing simulators can produce reads that resemble contemporary ONT data.

ONT simulators differ fundamentally in how they generate reads. Some tools rely on explicit training with real data, learning an empirical error model by aligning ONT reads to a reference genome and then using it to introduce errors into new sequences ^16,17,19^. Others use pre-trained models that encapsulate typical error behaviour for a given technology and chemistry. Others generate reads from parametric distributions specified by the user, without direct training on empirical data ^18,20^. These design choices affect not only how realistic the resulting reads are for a given technology snapshot, but also how easily the simulator can be adapted as ONT chemistries and basecalling models evolve. In particular, simulators that train an error profile from alignments of real reads to a reference have the potential to adapt to new read types.

In this context, ‘realism’ refers to how closely simulated reads resemble real ONT data across multiple dimensions that matter for downstream analysis. These include basic read-level metrics, such as the distributions of read length, read accuracy and FASTQ Q-scores, together with relationships between these metrics and finer-scale error profiles. The primary goal of this study is therefore to evaluate which currently available read simulators produce the most realistic data for modern ONT reads, focusing on faithfulness to empirical read characteristics.

## 2. Methods

### 2.1. Simulator selection

A literature search identified seven long-read simulators compatible with ONT reads. SimON-reads ^21^ was excluded because it failed to produce reads under any of the specified configuration flags and was deemed non-functional for this analysis. The other six simulators were successfully evaluated in this study: Badread v0.4.1 ^18^, LongISLND v0.9.5 (`c458f56`) ^19^, lrsim v0.7^22^, NanoSim v3.2.3 ^16^, PBSIM3 v3.0.5^17^, and SimLoRD v1.0.4 ^20^.

### 2.2. Reference data

Real ONT reads used as the empirical benchmark were generated from *Listeria innocua* strain AUSMDU00097349 (accession SAMN46906078) on an ONT R10.4.1 flow cell and basecalled with Dorado v1.1.1 ^23^ using sup@v5.2.0, the latest DNA basecalling model available for this chemistry at the time of analysis. The real read set comprised 680,482 reads with a total length of 3,803 Mbp, mean length of 5,588 bp and N50 length of 8,923 bp. The reads were split into two non-overlapping sets: a test set of 53,093 randomly selected reads (chosen to total 100*×* depth) and a training set containing the remaining 627,389 reads. The test set was used to evaluate real-read properties, while the training set was used to train simulators.

Simulated reads were generated from a complete, closed assembly of the same *L. innocua* strain. This 2,972,545 bp genome was produced in a previous study from the same ONT read set, with Illumina reads used for polishing ^12^. Because the assembly was carefully resolved and polished, it provided a reliable ground truth for both read simulation and read evaluation.

### 2.3. Simulator configuration

Each simulator was configured to maximise the realism of generated reads. For simulators with explicit training or model-building steps (Badread, LongISLND, lrsim and NanoSim), models were built from the real read training set, using alignments to the reference genome where required. PBSIM3 was run in its sampling mode, using real training reads as the source of read lengths and FASTQ quality strings, and SimLoRD was configured to sample read lengths from the training reads. For options controlling overall error rate, error variability or error-type ratios, we used values estimated from the training reads. All tools were set to generate read sets of 100*×* depth (∼300 Mbp). Exact commands are documented in Supplementary Methods (methods.md) and are publicly available in the accompanying GitHub repository.

### 2.4. Evaluation metrics

Simulated and real test reads were compared across multiple read-level and base-level metrics to assess realism. We did not rely on formal statistical tests, because with datasets of this size, even trivial differences can be statistically significant. Instead, we compared the shapes of distributions and the relationships between variables in simulated and real data, which provides a more direct and interpretable measure of realism.

Read lengths were extracted from FASTQ files and summarised using seqkit v2.13.0 ^24^. Read length distributions were visualised as histograms using the same bin widths and axis limits across all datasets to enable direct visual comparison.

Real and simulated reads were aligned to the reference genome using minimap2 v2.30 ^25^. Secondary alignments were removed, and identity was then calculated as the number of matching bases divided by the alignment length. Empirical read Q-scores were computed as *Q* = *−*10*×*log_10_(1*−*identity) for each read. Reported read Q-scores were calculated by converting each Phred-scaled quality value in the FASTQ file to an error probability, averaging those probabilities across the read and converting the mean error probability back to a Q-score.

To examine FASTQ Q-scores at the per-base level, each base was classified from the alignment as a match, mismatch or insertion, then grouped by its reported FASTQ Q-score. Because deletions have no corresponding read base and therefore no Q-score, they were represented indirectly by adjacent bases counted at half weight. For each reported Q-score, we then calculated the observed error rate from the counts of mismatches, insertions and deletion-associated bases, and used these values to generate calibration plots.

To characterise the error profiles of the real and simulated reads, we analysed two specific error types: 3-mer substitutions and homopolymer lengths. For 3-mer substitutions, we counted substitutions in which the middle base differed and the immediate flanking bases matched, for which there are 96 possibilities when reverse-complement equivalents are collapsed. The rate of each was expressed as the number of observed substitutions divided by the number of occurrences of the reference 3-mer in the aligned sequence. Homopolymer lengths of 3–10 bp were assessed using Pomoxis v0.3.16 ^26^, and we defined error rate as the fraction of homopolymers in reads with an incorrect length. These analyses were not intended to capture all aspects of ONT sequencing errors, but instead provide two targeted, interpretable and directly comparable measures of error realism.

## 3. Results

### 3.1. Simulator design and training strategies

The six evaluated read simulators (Badread, LongISLND, lrsim, NanoSim, PBSIM3 and SimLoRD) differ in how they model read length, sequence errors and Q-scores (Table 1) ^16–20,22^. Badread uses parametric read length and identity distributions, with trained *k* -mer error and Q-score models. LongISLND uses a non-parametric model that samples bases and Q-scores from training reads with matching extended-*k* -mer contexts. lrsim uses a model derived from alignment statistics for read lengths and indel-size structure, with user-specified overall error parameters. NanoSim fits read length, alignment, Q-score and error distributions from aligned reads, using Markov models for error structure. PBSIM3 was run in sample mode, where it copies read lengths and Q-score strings from real reads and uses a user-specified error-type ratio. SimLoRD is based on a PacBio circular consensus sequencing (CCS) pass-count model; in this study, we disabled CCS-style behaviour to better approximate ONT sequencing, sampled read lengths from the training reads and supplied error parameters manually.

**Table 1.**
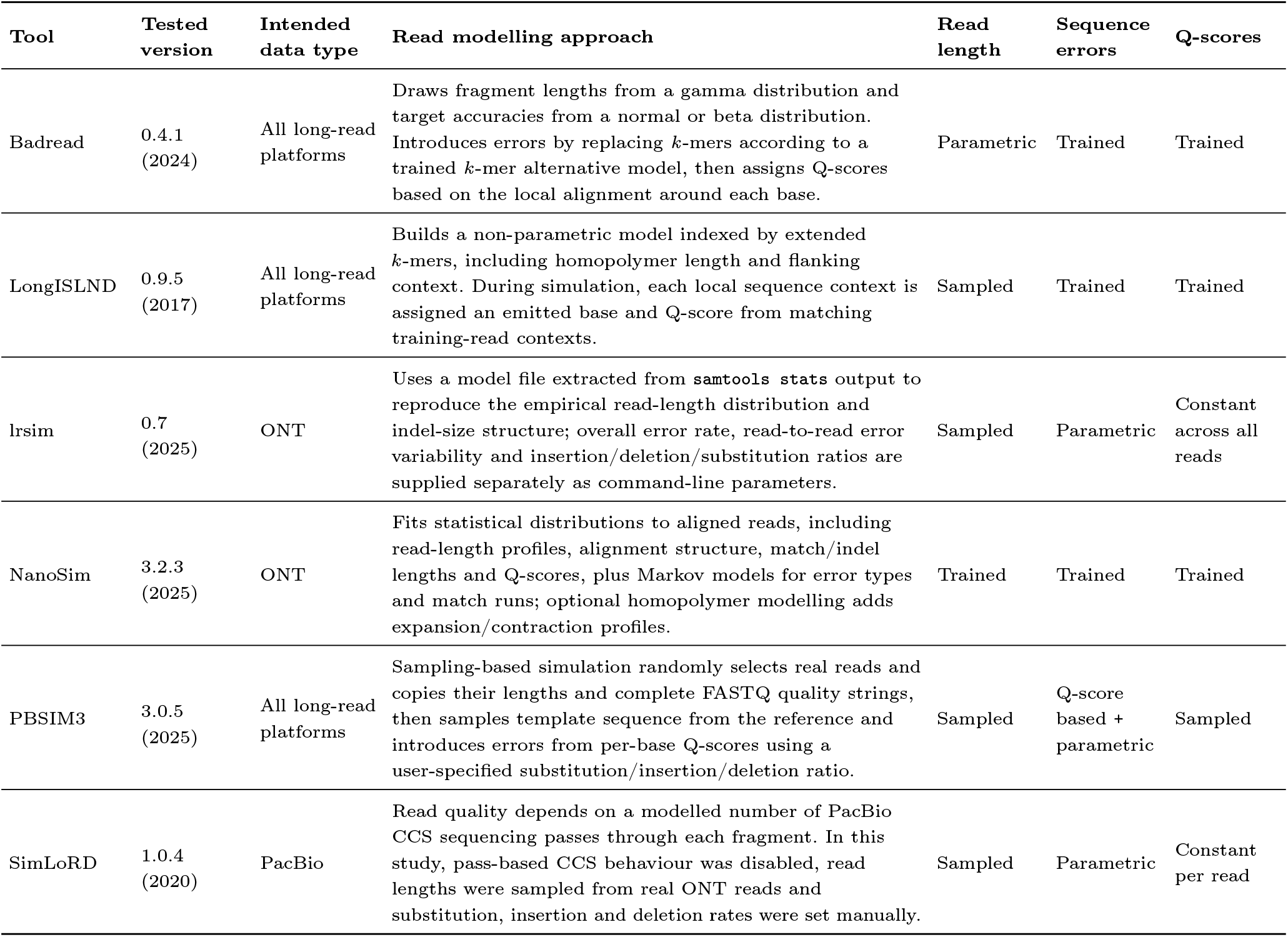
Key characteristics of the six evaluated ONT read simulators.

| Tool | Tested version | Intended data type | Read modelling approach | Read length | Sequence errors | Q-scores |
| --- | --- | --- | --- | --- | --- | --- |
| Badread | 0.4.1 (2024) | All long-read platforms | Draws fragment lengths from a gamma distribution and target accuracies from a normal or beta distribution. Introduces errors by replacing $k$-mers according to a trained $k$-mer alternative model, then assigns Q-scores based on the local alignment around each base. | Parametric | Trained | Trained |
| LongISLND | 0.9.5 (2017) | All long-read platforms | Builds a non-parametric model indexed by extended $k$-mers, including homopolymer length and flanking context. During simulation, each local sequence context is assigned an emitted base and Q-score from matching training-read contexts. | Sampled | Trained | Trained |
| lrsim | 0.7 (2025) | ONT | Uses a model file extracted from `samtools stats` output to reproduce the empirical read-length distribution and indel-size structure; overall error rate, read-to-read error variability and insertion/deletion/substitution ratios are supplied separately as command-line parameters. | Sampled | Parametric | Constant across all reads |
| NanoSim | 3.2.3 (2025) | ONT | Fits statistical distributions to aligned reads, including read-length profiles, alignment structure, match/indel lengths and Q-scores, plus Markov models for error types and match runs; optional homopolymer modelling adds expansion/contraction profiles. | Trained | Trained | Trained |
| PBSIM3 | 3.0.5 (2025) | All long-read platforms | Sampling-based simulation randomly selects real reads and copies their lengths and complete FASTQ quality strings, then samples template sequence from the reference and introduces errors from per-base Q-scores using a user-specified substitution/insertion/deletion ratio. | Sampled | Q-score based + parametric | Sampled |
| SimLoRD | 1.0.4 (2020) | PacBio | Read quality depends on a modelled number of PacBio CCS sequencing passes through each fragment. In this study, pass-based CCS behaviour was disabled, read lengths were sampled from real ONT reads and substitution, insertion and deletion rates were set manually. | Sampled | Parametric | Constant per read |

### 3.2. Read length distribution

LongISLND, lrsim, NanoSim, PBSIM3 and SimLoRD all used the training reads to model or sample read lengths, allowing them to approximate the empirical ONT read length distribution. Therefore, their simulated read length distributions were similar to the real reads (Table S1 and Figure S1).

In contrast, Badread generates read lengths from a gamma distribution defined by mean read length and standard deviation. It was therefore the only tool which did not closely replicate the empirical distribution.

### 3.3. Empirical and reported Q-score distributions

Each read that could be aligned to the reference genome was assigned an empirical Q-score, indicating its true accuracy. Each read also had a reported Q-score calculated from its FASTQ quality values. Comparing the joint and marginal distributions of these two Q-scores therefore provided a direct visual measure of how closely each simulated dataset resembled the real reads.

Only PBSIM3 was able to closely reproduce the signature of the real reads, including their distinctive bimodal distribution with a lower peak at *∼*Q8 and a higher peak at *∼*Q23 (Figure 1). This likely reflects PBSIM3’s sampling-based strategy: it copies read lengths and per-base FASTQ quality strings from training reads, samples template sequences of matching length from the reference genome, and introduces errors according to the sampled Q-scores. PBSIM3’s reads only departed from the real reads in a few subtle ways: the real read distributions extended to lower qualities than the PBSIM3 reads, and the real reads had more outliers where empirical and reported Q-scores were in stark disagreement (e.g. reported Q20 but empirical Q10).

**Figure 1:**
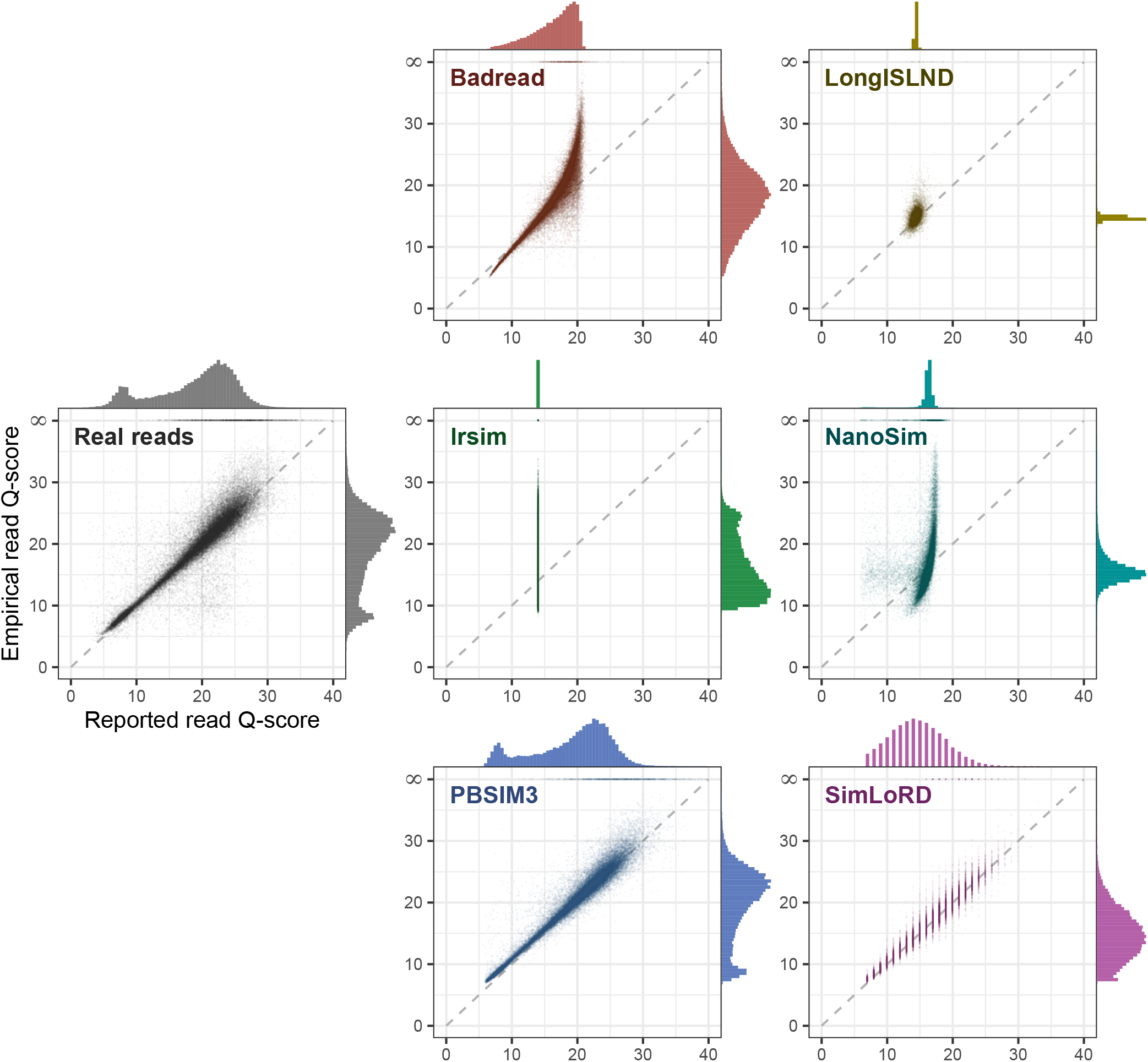
Empirical and reported read Q-scores. In each panel, the central scatter plot shows each aligned read’s reported Q-score (x-axis) and empirical Q-score (y-axis). The reported read Q-score was calculated from the FASTQ quality string by converting each base Q-score to an error probability, averaging those probabilities across the read, and converting the mean error probability back to a Q-score. The empirical read Q-score was calculated from the read-to-reference alignment identity, converted to Q-score. The top y-axis tick is labelled ∞ because some reads contained no errors, giving an empirical error rate of zero and therefore an infinite empirical Q-score. The top marginal histogram for each panel shows the distribution of reported read Q-scores, and the right marginal histogram shows the distribution of empirical read Q-scores. Histogram bars are weighted by read length, so the marginal axes represent the number of bases in each bin. A simulator is more realistic if its plot closely matches that of the real reads.

All other tools departed from the real-read Q-score distributions. Badread had a wide spread of empirical Q-scores, but its reported Q-scores rarely exceeded Q20. LongISLND produced very narrow distributions, with most reads in the Q14–15 range. lrsim produced a range of empirical Q-scores but did not model reported Q-scores at all, using the same Phred score character for every base in every read. NanoSim produced narrow distributions for both empirical and reported Q-scores. SimLoRD produced wide distributions, but it used only a single Phred score character in each read, leading to a discrete reported Q-score distribution.

When Q-scores were analysed at the per-base level rather than the whole-read level, PBSIM3 was again the most realistic simulator. Badread, LongISLND and PBSIM3 all broadly reproduced the distribution of reported base Q-scores for correctly called bases (Figure S2), but only PBSIM3 also closely matched the distribution of reported base Q-scores for error-associated bases (Figure S3) and for the Q-score calibration curve (Figure S4).

### 3.4. Error profiles

Errors in the real reads were strongly context dependent: the 96 3-mer substitution error rates spanned nearly two orders of magnitude, from 0.0263% to 2.01% (Figure 2). This is consistent with ONT sequencing’s raw-current signal (the “squiggle”), in which some *k* -mers produce less distinct signals and are therefore more prone to basecalling errors ^27^. Badread and LongISLND were the only simulators to reproduce this broad pattern, although neither matched the real profile exactly. In contrast, lrsim, NanoSim, PBSIM3 and SimLoRD assigned roughly uniform error rates to all 3-mer substitution types, failing to reproduce the context specificity seen in the real reads.

**Figure 2:**
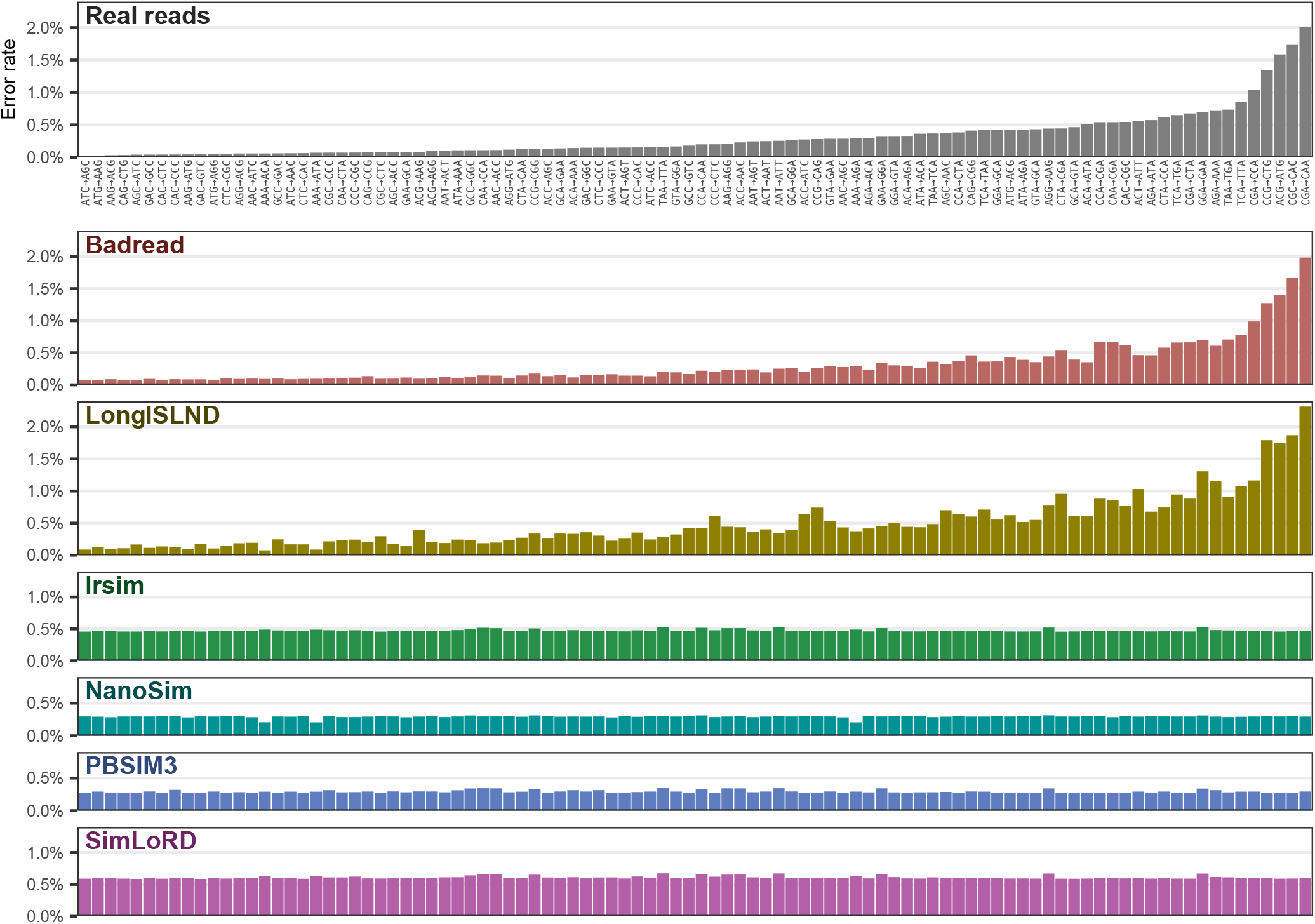
Substitution rates by 3-mer context. Error rates for the 96 distinct 3-mer substitution types, in which only the middle base changes and reverse-complement equivalents are combined. Substitutions are ordered from lowest to highest error rate in the real reads, and the same order is used for all simulated datasets. A simulator is more realistic if its plot closely matches that of the real reads.

Homopolymer error rates in the real ONT reads increased with homopolymer length, with the rise becoming steeper from 8 bp onward (Figure 3). LongISLND reproduced this overall pattern most closely, although its curve showed an unusual drop from 4-mer to 5-mer homopolymers. NanoSim produced a markedly different profile, with much lower error rates for homopolymers of 5 bp and longer despite being run with explicit homopolymer-length modelling for those lengths, showing that this feature did not reproduce the empirical pattern well. Badread, lrsim, PBSIM3 and SimLoRD generally showed increasing error with homopolymer length, but none followed the real-read curve closely, particularly for longer homopolymers.

**Figure 3:**
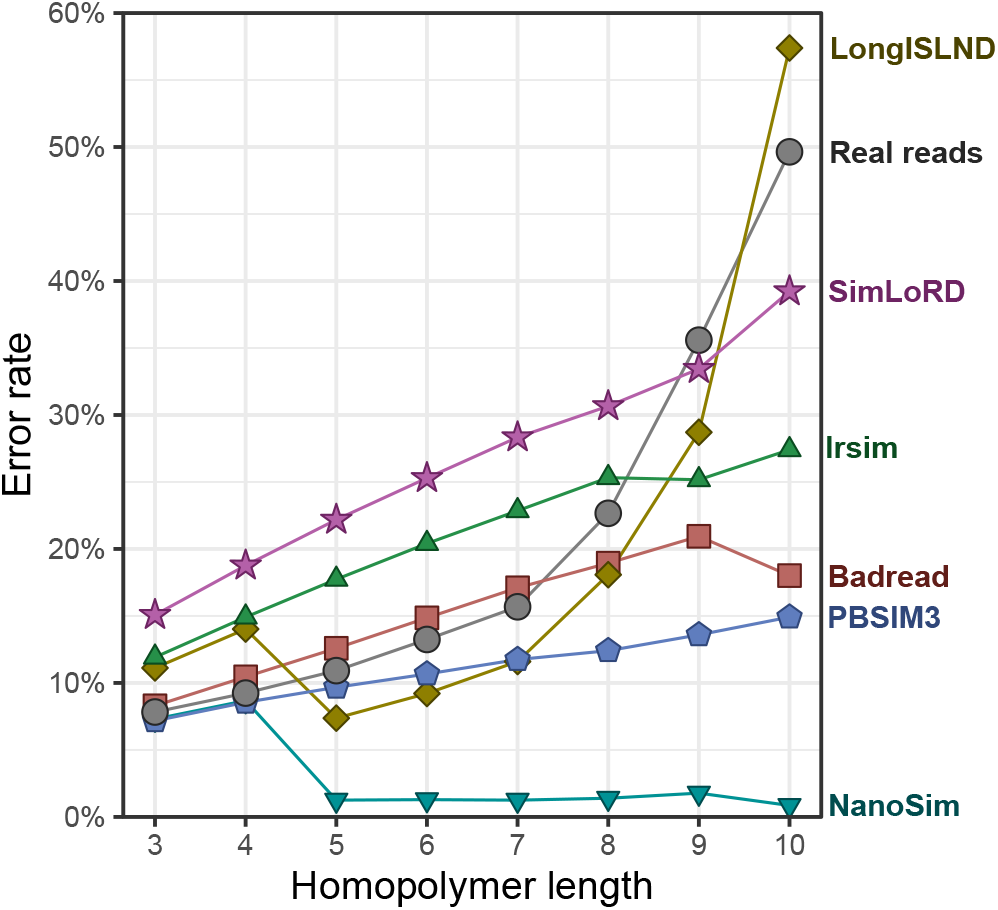
Homopolymer error rates. Homopolymer error rate as reported by Pomoxis, plotted as a function of homopolymer length for real reads and six ONT read simulators. Error rate is defined here as the fraction of reads in which the homopolymer length was not represented correctly. A simulator is more realistic if its curve closely matches that of the real reads.

## 4. Discussion

This study benchmarked six ONT-compatible read simulators against contemporary R10.4.1 ONT real reads, with the goal of identifying which tools best reproduce empirical read characteristics. That is, we judged simulators by how closely their outputs matched real reads across features relevant to downstream use, such as method development and pipeline validation. Our results demonstrate that no simulator perfectly captures all aspects of modern ONT reads, but PBSIM3, Badread and LongISLND performed best, depending on the key metric considered.

PBSIM3 performed strongly overall and was the most realistic simulator for several key properties of the read sets. Its simulated reads closely matched the real data in read length, empirical read accuracy and FASTQ Q-scores (Figure 1), making it a practical choice for users who want simulated datasets that resemble real ONT whole-genome sequencing output in broad terms. This realism likely comes from its sampling-based approach, where quality strings are copied from empirical data rather than generated de novo. However, a limitation of PBSIM3 was that it was less successful at reproducing the error profile of the real data (Figures 2 and 3).

Badread offers users parametric control over the simulated reads, which can be valuable when specific read-set properties are needed. This flexibility came at the expense of realism: its read length and accuracy distributions deviated from those of the real reads (Figures S1 and 1). Badread modelled substitution errors well (Figure 2) but captured homopolymer-length errors less well (Figure 3). Further, as Badread is implemented entirely in Python, it was slower to run than other tools.

LongISLND performed best at error modelling, capturing substitutions and handling homopolymer-length errors reasonably well (Figures 2 and 3). However, its distributions of empirical and reported read Q-scores were unrealistically narrow, limiting its potential use as a simulator (Figure 1).

The remaining simulators were less suitable for realistic ONT read simulation. lrsim and SimLoRD were limited by their treatment of FASTQ Q-scores: lrsim assigns the same Q-score to every base in the entire output, while SimLoRD assigns a single Q-score per read. This makes both tools unsuitable for applications which use per-base Q-scores. NanoSim produced an unrealistically narrow distribution of read-level accuracy (Figure 1), an unrealistically broad distribution of base-level accuracy (Figures S2 and S3) and poor modelling of homopolymer-length errors, despite being explicitly configured to simulate them.

In this study, we did not exhaustively explore every possible configuration of each tool. We aimed to run each one in the way that we judged most likely to maximise overall realism. Other configurations may have produced different results. For example, we enabled NanoSim’s homopolymer modelling because it is intended to better reflect ONT data, but it is possible that NanoSim would have performed better on our dataset with this feature disabled. PBSIM3 includes an error-model HMM mode that may improve error realism, but this comes with trade-offs: read lengths and Q-scores are then generated from parametric models rather than sampled from empirical data, and it does not support training a custom model on user-provided data.

Another limitation is that our benchmarking focused only on WGS. Other applications of long-read sequencing, such as transcriptomics, amplicons and metagenomics, may prioritise different properties of simulated reads. We also did not assess how reads were distributed across the reference genome. Real sequencing datasets may show positional biases in read start sites or coverage, arising from library preparation, sequence composition or other experimental factors, and these biases could be important for some downstream analyses.

Although this benchmark used a bacterial genome and ONT whole-genome sequencing data, we expect the conclusions to generalise more broadly. The bacterial dataset was a practical choice because the read sets were not too large and the reference genome was highly accurate, but the properties we assessed are not specific to bacterial genomics. We therefore expect similar conclusions to hold for other organisms, provided that simulations are trained on matching empirical data, such as real human reads for human-read simulation. We also note that this study considered only ONT data, and other long-read platforms, such as PacBio HiFi, have different error profiles. Nevertheless, given the underlying approaches used by the tools, we suspect their relative strengths and weaknesses would remain similar on data from other platforms. Future work could directly test how well these conclusions generalise beyond ONT sequencing.

## 5. Conclusions

No simulator produced reads that looked realistic across all tested metrics. When used in its sampling-based mode, PBSIM3 is likely the best choice for most users. It was easy to run and performed well on several key properties of ONT whole-genome sequencing data, particularly read length and read-level accuracy metrics, although it failed to model error types well. Badread did better at modelling some error types and offers parametric control over the simulated read set, but at the cost of less realism and substantially slower run times. LongISLND most closely matched the tested error features, but its simulated reads had an unrealistically narrow accuracy distribution. The remaining simulators each showed limitations that make them less suitable for general use. More broadly, the fact that no tool performed well on all metrics highlights an opportunity for improved long-read simulators that can combine realistic read-level distributions with more faithful modelling of sequencing errors.

## 6. Availability of data and materials

Detailed commands, custom scripts and complete analysis outputs are publicly available at: github.com/mtaouk/ONT-Read-Simulator-Benchmarking.

The reference genome and full read sets used in this study are available at: doi.org/10.26188/32194977.

## 9. Authors’ contributions

R.R.W. conceived the study. M.L.T. and R.R.W. performed analyses and drafted the manuscript. D.J.I. and R.R.W. supervised the work, contributed to study design and interpretation of results, and revised the manuscript critically. All authors read and approved the final manuscript.

## Supplementary Information

Supplementary methods are available at: github.com/mtaouk/ONT-Read-Simulator-Benchmarking.

**Table S1.** Read length statistics.

| Dataset | Number of reads | Total bases (bp) | Min length (bp) | Max length (bp) | Mean length (bp) | N50 length (bp) |
| --- | --- | --- | --- | --- | --- | --- |
| Real reads | 53,093 | 297,265,978 | 5 | 826,029 | 5,599.0 | 8,994 |
| Badread | 53,256 | 297,257,226 | 1 | 79,424 | 5,581.7 | 10,509 |
| LongISLND | 53,956 | 297,851,844 | 37 | 181,950 | 5,520.3 | 8,758 |
| lrsim | 54,022 | 297,723,160 | 3 | 79,543 | 5,511.1 | 8,784 |
| NanoSim | 53,216 | 317,425,632 | 50 | 126,188 | 5,964.9 | 8,988 |
| PBSIM3 | 53,541 | 297,262,126 | 91 | 89,449 | 5,552.0 | 8,785 |
| SimLoRD | 53,202 | 299,722,584 | 5 | 369,970 | 5,633.7 | 8,995 |

**Figure S1:**
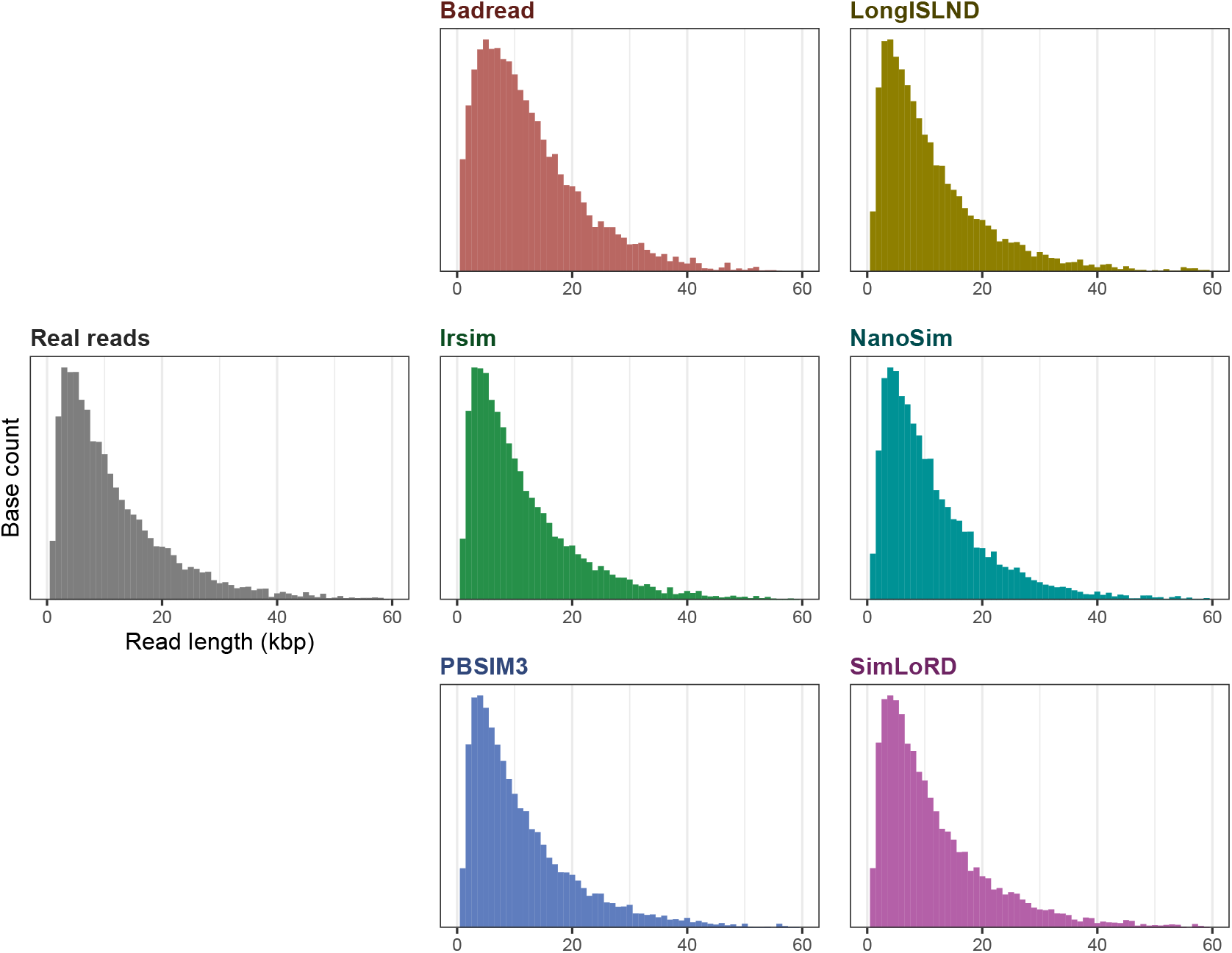
Read length distributions. Histograms are shown for the real and simulated datasets. The x-axis shows read length in 1 kbp bins. Histogram bars are weighted by read length, so the y-axis represents the total number of bases contributed by reads in each length bin rather than the number of reads. A simulator is more realistic if its histogram closely matches that of the real reads.

**Figure S2:**
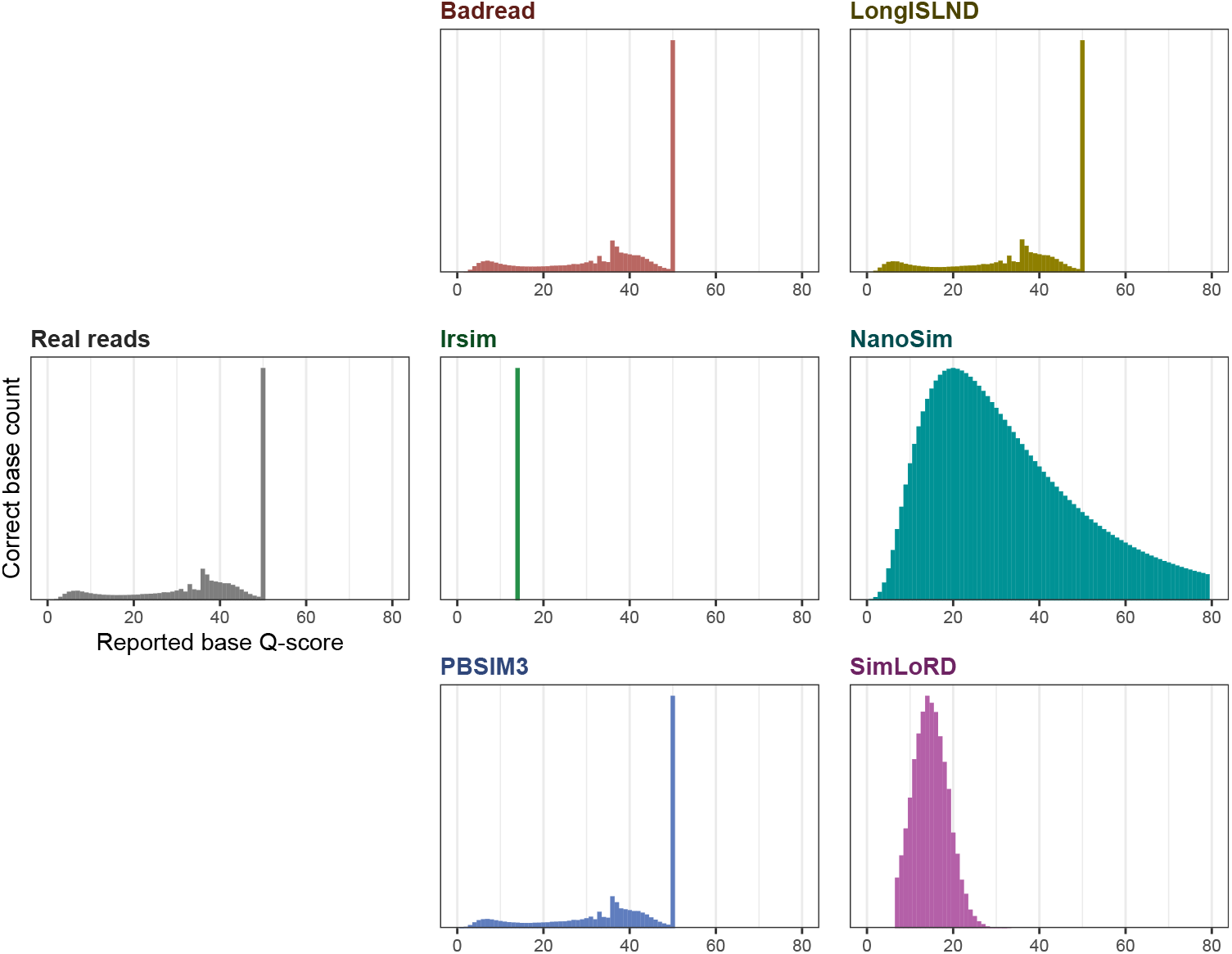
Correct base Q-score distribution. Reported base Q-score distributions for correctly called bases in the real and simulated datasets. In each panel, bars show the number of bases in each reported base Q-score bin. A base was counted as correct only if it matched the reference and was not adjacent to a deletion in the alignment. A simulator is more realistic if its histogram closely matches that of the real reads.

**Figure S3:**
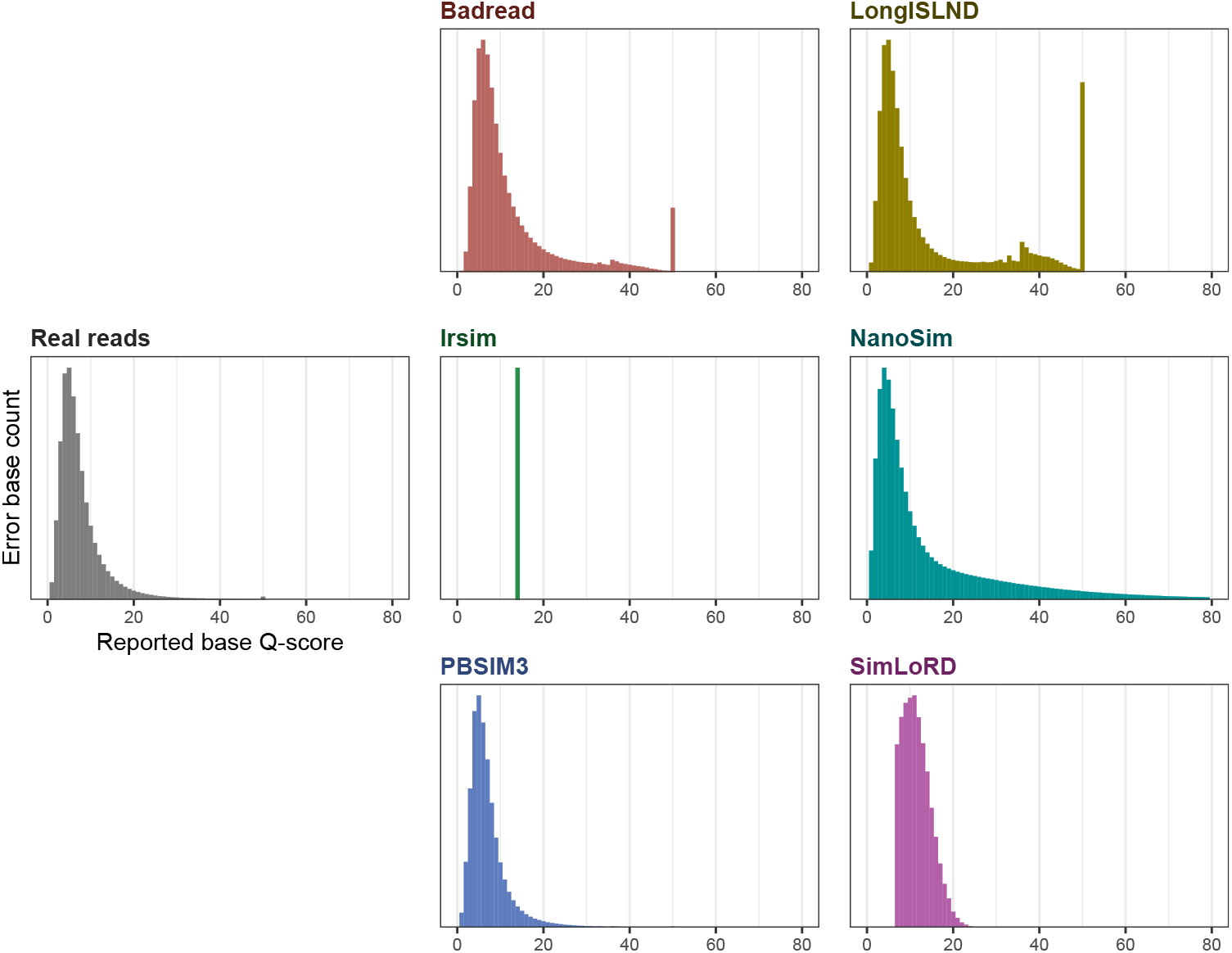
Error-associated base Q-score distribution. Reported base Q-score distributions for error-associated bases in the real and simulated datasets. In each panel, bars show the number of error-associated bases in each reported base Q-score bin. Error-associated bases were defined as mismatches and insertions, plus deletion-associated events estimated as half the number of matched bases adjacent to deletions, because deleted bases themselves have no reported Q-score. A simulator is more realistic if its histogram closely matches that of the real reads.

**Figure S4:**
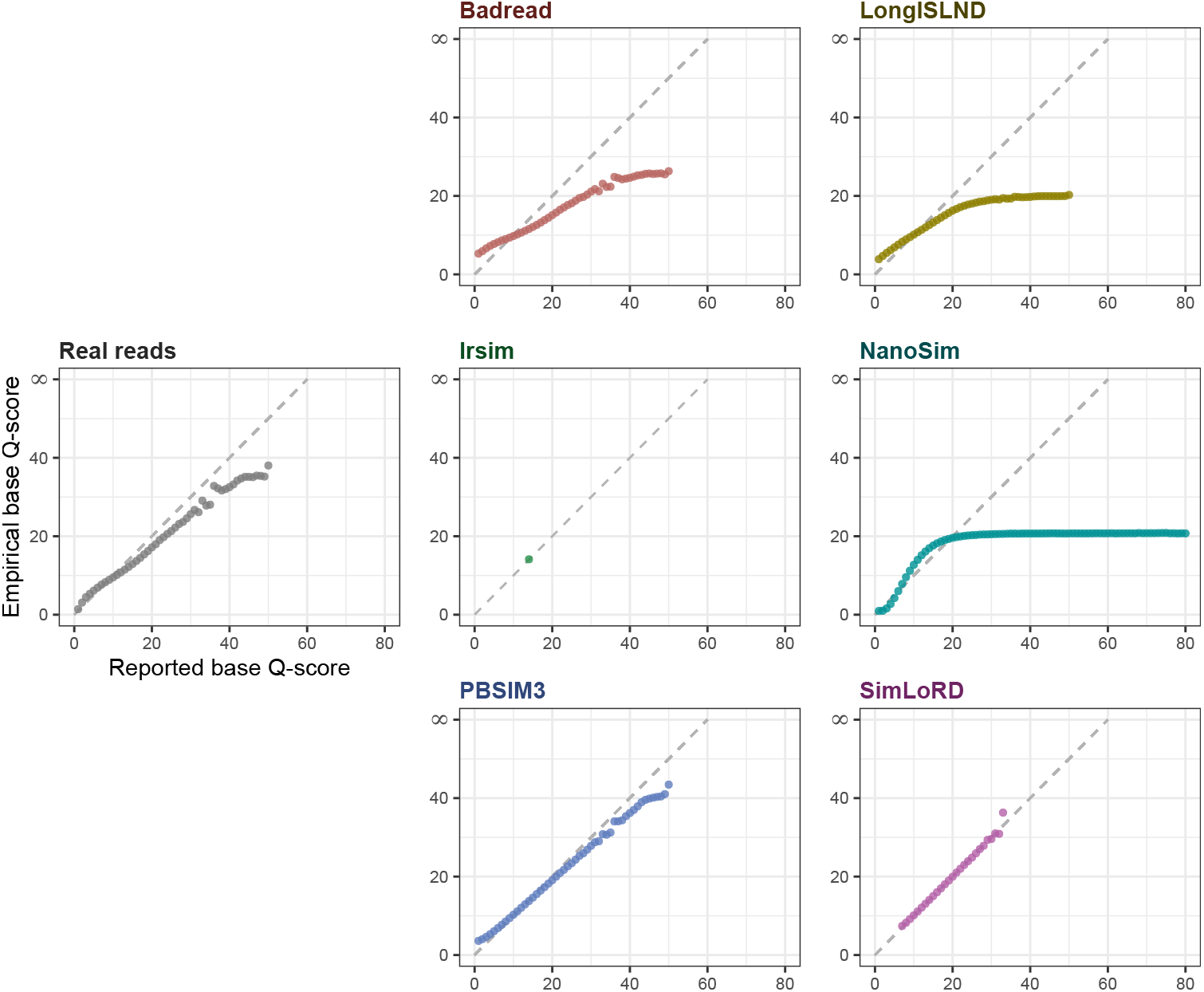
Q-score calibration curve. Per-base Q-score calibration for real and simulated datasets. Each point represents one reported base Q-score bin (x-axis), and its y-value is the empirical base Q-score calculated from the observed error rate among bases with that reported Q-score. Empirical error rate was defined as the fraction of bases in the bin that were mismatches or insertions, plus deletion-associated events estimated as half the count of matched bases adjacent to deletions. The dashed diagonal indicates perfect calibration, where reported and empirical base Q-scores are equal.

